# ISMI-VAE: A Deep Learning Model for Classifying Disease Cells Using Gene Expression and SNV Data

**DOI:** 10.1101/2023.07.28.550985

**Authors:** Han Li, Ying Wang, Yongxuan Lai, Feng Zeng, Fan Yang

## Abstract

Various studies have linked several diseases, including cancer and Covid-19, to single nucleotide variations (SNV). Although scRNA-seq technology can provide SNV and gene expression data, few studies have integrated and analyzed these multimodal data. To address this issue, this paper introduces Interpretable Single-cell Multimodal Data Integration Based on Variational Autoencoder (ISMI-VAE). ISMI-VAE leverages latent variable models that utilize the characteristics of SNV and gene expression data to overcome high noise levels, and uses deep learning techniques to integrate multimodal information, map them to a low-dimensional space, and classify disease cells. Moreover, ISMI-VAE introduces an attention mechanism to reflect feature importance and analyze genetic features that could potentially cause disease. Experimental results on three cancer data sets and one Covid-19 data set demonstrate that ISMI-VAE surpasses the baseline method in terms of both effectiveness and interpretability, and can effectively identify disease-causing gene features.

## 1 Introduction

With the development of sequencing technology, we are now able to easily access various modal data from the same cells. These modal data include gene expression, DNA methylation, copy number variation, etc[1, 2]. Obviously, the data of different modalities do not contain exactly the same information. Compared with single-modal data, multimodal data can better reflect the state of cells and thus provide more accurate analysis for biological research. With the emphasis on multimodal data, many multimodal integration analysis methods have been proposed[3–5]. The objectives of these methods include clustering of multimodal data, prediction of diseases, and analysis of inter-modal relationships. These multimodal analysis methods have achieved better results and provided deeper biological insights about cells compared to single-modal analysis methods.

The scRNA-seq data is commonly used, and many multimodal analysis methods utilize it to generate gene expression data for clustering or classification tasks. However, scRNA-seq can be used not only to provide gene expression data, but also to generate SNV data[6]. Unlike gene expression data that count reads on different genes, SNV data count mutations at different loci. SNV data record variant and reference read counts at different loci, and obtain genotype information of the locus based on the ratio of variant to reference read counts. SNV plays an important role in the human body, not only affecting development and appearance, but also influencing the development of many diseases, such as Alzheimer’s disease and cancer[7, 8]. Because of this role, SNV data have been used in many research areas, including eQTL[9], demultiplexing[10], etc.

scRNA-seq is able to provide both single-cell-scale gene expression data and SNV data. With scRNA-seq data alone, we are able to perform integration of multimodal data and improve the effectiveness of the task. However, few such studies have been conducted yet. In fact there is a great potential to do so. On the one hand, gene expression data can indirectly reflect RNA-based protein expression levels, but cannot reflect gene sequence information, which SNV happens to reflect[11]. On the other hand, many diseases not only affect gene expression data, but more importantly, it is influenced by SNVs or even caused by SNVs[12, 13]. Therefore, combining gene expression data and SNV data can better reveal the mechanism of action of diseases. In this paper, we propose Interpretable Single-cell Multimodal Data Integration Based on Variational Autoencoders(ISMI-VAE), to solve the integration problem of gene expression data and SNV data. We constructed a latent variable model based on the distribution of these two data and designed a neural network model based on variational autoencoders[14]. Considering the important role of SNV data in the disease process, we predict the disease cells by fusion of multimodal data. In addition, we additionally considered the importance of interpretability and proposed an attention modality to explain the importance of features, which was used to analyze the disease-causing genes.

The contributions of our work can be summarized as follows:

1. We proposed the model ISMI-VAE for integrating SNV data and gene expression data and classifying disease cells.
2. ISMI-VAE constructs a latent variable model to describe the distribution of data and uses a neural network model to integrate multimodal data into a low-dimensional feature space and to predict disease cells.
3. ISMI-VAE proposes an attention module that uses the weights of the attention vector to reflect the importance of gene features as a way to determine genes or SNVs that are highly associated with disease.
4. Experiments demonstrate that integrating SNV data can effectively improve the classification of the model, and ISMI-VAE has good performance in integrating multimodal data as well as interpretability analysis.

## 2 Results

## 3 Overview of ISMI-VAE

The basic idea of ISMI-VAE is to combine attention mechanism and variational autoencoder(Fig. 1). We utilize variational autoencoders to fit the distribution of gene expression data and SNV data, and map the integrated data into the latent variable space, and then classify disease cells. We use the attention mechanism to assign an attention weight to each feature, and analyze the feature importance according to this weight.

**Fig. 1.**
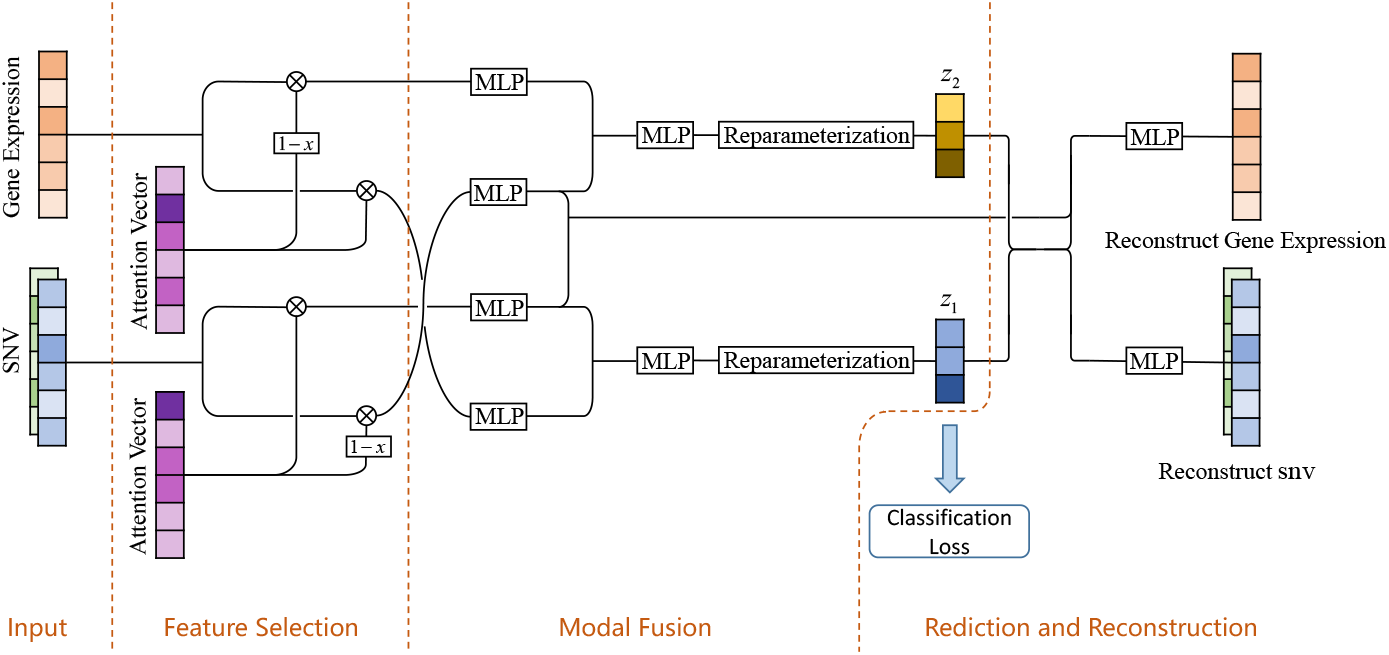
overview of the ISMI-VAE. Input data include gene expression data and SNV data. The model is able to be divided into three parts. In the feature selection part, the input data are divided into significant and non-significant features by the attention vector. In the modality fusion part, the data of different modalities are dimensionally reduced and fused by multiple multi-layer perceptrons. Meanwhile, the important features generate a latent variable for predicting the labels and reconstructing the data, and the non-important features generate a latent variable for reconstructing the data only. These tasks are completed in the final rediction and reconstruct part.

The neural network model of ISMI-VAE can be divided into three parts. First, ISMI-VAE processes the data of each modality separately. For a given data, ISMI-VAE will multiply it with a randomly initialized attention vector to obtain a feature vector highly correlated with the disease. In this process, the obtained feature vector miss part of the original data, which makes it difficult for the model to reconstruct the data in the end. To avoid this, ISMI-VAE retains missing data. In the second part, ISMI-VAE integrates data from each model and generates latent variables. Specifically, the input data is first dimensionally reduced through a multi-layer perceptron. Then, data from different modalities will be integrated through concatenation. Finally, another multilayer perceptron will further dimensionally reduce the concatenated vectors and use them to generate latent variables. The disease-related feature vectors and missing data obtained in the first part will follow the same steps to generate latent variables, z1 and z2, respectively. In the third part, z1 is used to predict diseased cells, and z1 and z2 are used to reconstruct the data.

## 4 Experiment settings

In this paper, we conducted four experiments. First, to evaluate the effectiveness of SNV data on disease cell classification and the multimodal information integration performance of ISMI-VAE, we conducted a multimodal data integration experiment. In this experiment, we evaluated and compared and the classification ability of the model under single-modal data and under multimodal data. After that, we compare the classification performance of ISMI-VAE with other multimodal integration models. We conducted experiments on 4 datasets, including 2 datasets used for classification of cancer types, one dataset used for classification between cancer and healthy cells and one dataset used for classification of the severity of Covid-19. After that, we performed a test of interpretability ability. We performed cell classification experiments with the key features found by ISMI-VAE and specifically analyzed the important features found on the Covid-19 dataset. Finally, we show the results of the parametric sensitivity analysis of ISMI-VAE.

The baseline methods we compared can be divided into three categories. The first category is single-modal traditional methods, including XGBoost[15] and Random Forest. The second category is multimodal traditional classification methods, including SMSPL[16] and DIABLO[17]. the third category is multimodal deep learning classification methods, including MOGONET[18] and MOMA[19].

The datasets we used for the experiments can be divided into two categories. First, SNVs are one of the important causes of cancer, and it is valuable to use SNVs to predict cancer types and early cancer occurrence, or to analyze key SNVs that cause cancer, so the first task of the experiment is cancer prediction. In this paper, three cancer datasets were selected for the experiments, named Patel, Pan-cancer, and Lung datasets.

The Patel dataset[20] contains five glioblastoma samples and two glioma samples from different patients, and after sequencing, a total of 862 cells data were obtained. We divided each cell population into training set and test set according to 1:1, and obtained 431 cells in the training set and 431 cells in the test set.

The Pan-cancer dataset[21] is a pan-cancer dataset, containing 17 cancer samples including skin cancer, liver cancer, breast cancer, etc. After sequencing, a total of 6157 cells are obtained. We also divide each cancer sample into training set and test set according to 1:1, then we get 3074 cells in training set and 3083 cells in test set.

The Lung dataset[22] contains 3 lung adenocarcinoma samples at IA stage and 3 healthy samples. 2000 cells were taken from each sample. Two lung cancer samples and two healthy samples were used as the training set and the remaining samples were used as the test set. There are 8000 cells in the training set and 4000 cells in the test set.

Several studies have shown that the severity of Covid-19 is also influenced by SNV[12], so the second task of the experiments is the prediction of Covid-19 severity. In this paper, a publicly available Covid-19 dataset was selected for the experiment.

The Covid-19 dataset[23] contains PBMC samples from six patients with severe Covid-19 and four patients with mild Covid-19. The data from three patients with severe Covid-19 and two patients with mild Covid-19 are used as the training set and the rest as the test set. There are 2897 cells in the training set and 2922 cells in the test set.

### 4.1 Data Progress

We used different methods to pre-process SNV data and gene expression data. For the gene expression data, we processed the data in the following 4 steps. In the first step, the genes with zero total expression are filtered out from the data. In the second step, the top 10,000 genes with the highest total expression level were selected as features. In the third step, the gene expression value are calculated based on log(*x/*10 + 1). Finally, each gene expression value was subtracted from the average value of that gene expressed on each cell. For the SNV data, the features with less than 10 non-zero expressed cells in either the alternate matrix and the reference matrix were filtered out first. The two matrices are then summed and the features that still have fewer than 200 non-zero values expressing cells are filtered out. After SNV data preprocessing, 61551 features remained in the Patel dataset, 6702 features in the Pan-cancer dataset, 21439 features in the Lung dataset, and 31289 features in the Covid-19 dataset.

### 4.2 ISMI-VAE can improve performance by combining SNV data and gene expression data

First, we verify that ISMI-VAE can make good use of multimodal information, and that SNV data can significantly enhance the classification effect of the model. In our experiments, we use multimodal data and single-modal data as model inputs respectively, and then compare the classification performance to observe whether the classification results of multimodal data outperform those of single-modal data. To obtain and measure the classification results of single-modal data, in this paper, we split ISMI-VAE into two parts, one for classified SNV data and the other for classified gene expression data. Specifically, the information fusion module in ISMI-VAE is removed, and the data of each modality are independently input into the feature selection module and the prediction and reconstruction module.

The experimental results are shown in Fig. 2. Comparing the classification results of gene expression data and SNV data, it can be found that the classifier can also have good classification performance by relying only on single-modal SNV data. This indicates that the SNV data contains more information that can help us identify the disease. In addition, on the Covid-19 dataset, the prediction using SNV is better than that using gene expression data, which may be because mutations affecting disease susceptibility are less likely to be reflected in gene expression. In contrast, on the cancer data, cancer-related mutations lead to large changes in gene expression; therefore, both SNV data and gene expression data are better able to distinguish cancer cells from normal cells.

**Fig. 2.**
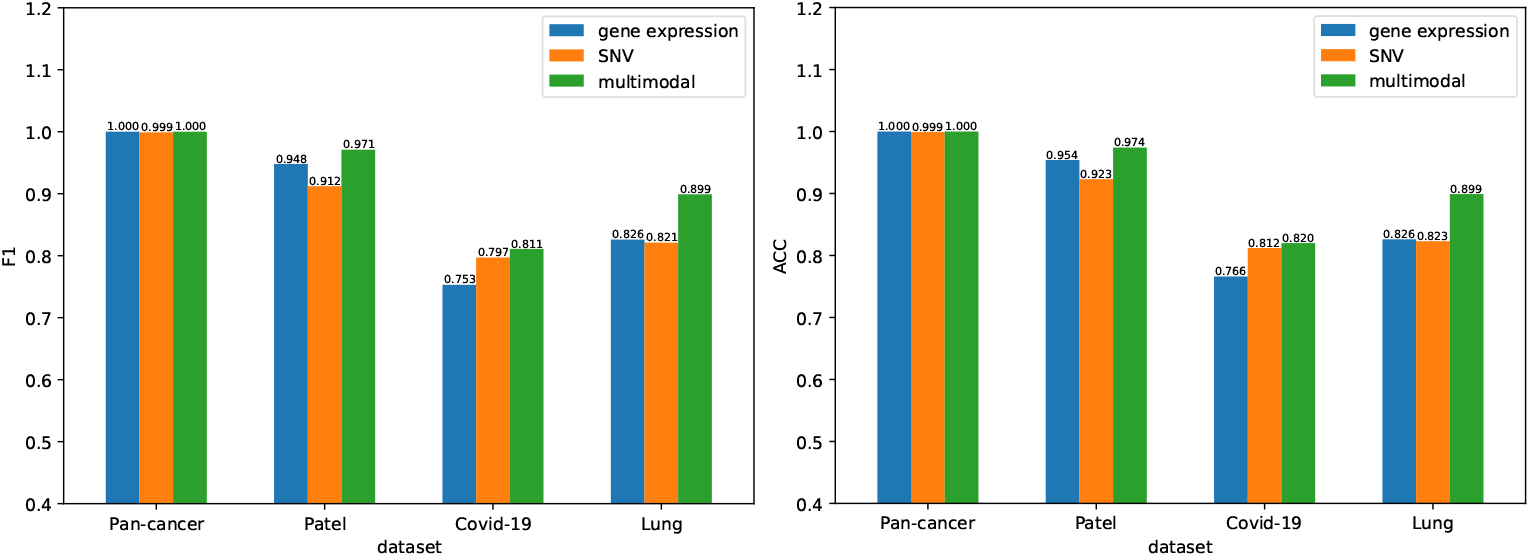
Classification effects on four datasets, where different colors represent gene expression data, SNV data, and multimodal data containing these two data as inputs to the model.

Comparing the classification performance of multimodal data and single-modal data shows that the classifier performance combining data from both modalities maintains the best results on all data, especially with the Lung dataset, the classifier performance was improved by about 6% for multimodal compared to gene expression data that performed better in single-modal data. This implies that SNV data contains information that is not present in gene expression data, and combining SNV and gene expression data can effectively improve the prediction of diseases.

### 4.3 ISMI-VAE has good disease prediction performance

We next evaluate the disease prediction ability of the model by classification effect. In our experiment, we use gene expression data and SNV data as input, and XGBoost, random forest, SMSPL, DIABLO, MOGONET and MOMA as baseline methods. Among them, XGBoost and Random Forest algorithms can only handle single-modal data, so we directly concat the data of two modalities as the input of these two algorithms. The four baseline methods, SMSPL, DIABLO, MOGONET, and MOMA, can only handle multimodal data in matrix form, but not SNV data, which is a special form of data, so in this experiment, we concat two matrices of SNV data together as the input of these algorithms.

The experimental results are shown in Table 1. In terms of the final classification results, the same batch prediction was overall better than the cross-batch prediction, and among these, the Pan-cancer dataset was the easiest to predict, with all methods except DIABLO reaching ACC and F1 metrics above 0.9. The baseline methods have their own advantages and disadvantages on these four datasets, for example, MOMA outperforms the other baseline methods on the Covid-19 and Lung datasets, while MOGONET achieves good results on the Patel dataset, outperforming the other baseline methods, but performs poorly on the Covid-19 dataset. The overall performance of SMSPL is average, but it slightly outperforms the other baseline methods on the Pan-cancer dataset. In general, the baseline method does not have consistent results on all data sets. However, ISMI-VAE outperformed all baseline methods on all four data. Among them, ISMI-VAE has a small performance improvement compared to the baseline method on the Pan-cancer dataset and Covid-19 dataset, with ACC improving by 0.9% and F1 improving by 0.9% and 1.1%, respectively. It has a larger performance improvement over the baseline method on the Patel and Lung datasets, with ACC improving by 3.5% and 2.4%, respectively, and F1 improving by 3.4% and 2.4%, respectively. It can also be seen that only ISMI-VAE predicts completely accurate on the Pan-cancer dataset, which indicates that ISMI-VAE not only has a good classification effect but also has good robustness to adapt to different datasets.

**Table 1.** Classification performance of ISMI-VAE and baseline methods.

| Dataset<br>Metrics | Pan-cancer |  | Patel |  | Covid-19 |  | Lung |  |
| --- | --- | --- | --- | --- | --- | --- | --- | --- |
|  | ACC | F1 | ACC | F1 | ACC | F1 | ACC | F1 |
| XGBoost | 0.919 | 0.925 | 0.883 | 0.889 | 0.696 | 0.729 | 0.798 | 0.8 |
| RF | 0.961 | 0.983 | 0.873 | 0.884 | 0.648 | 0.709 | 0.836 | 0.837 |
| SMSPL | 0.991 | 0.991 | 0.921 | 0.911 | 0.594 | 0.373 | 0.796 | 0.789 |
| DIABLO | 0.812 | 0.823 | 0.847 | 0.854 | 0.732 | 0.7 | 0.64 | 0.606 |
| MOMA | 0.982 | 0.966 | 0.893 | 0.881 | 0.811 | 0.8 | 0.875 | 0.875 |
| MOGONET | 0.99 | 0.975 | 0.93 | 0.925 | 0.734 | 0.76 | 0.873 | 0.874 |
| Ours | **1** | **1** | **0.965** | **0.959** | **0.82** | **0.811** | **0.899** | **0.899** |

We also show the ROC curves and AUC of each method. the experimental results are shown in Fig. 3.

**Fig. 3.**
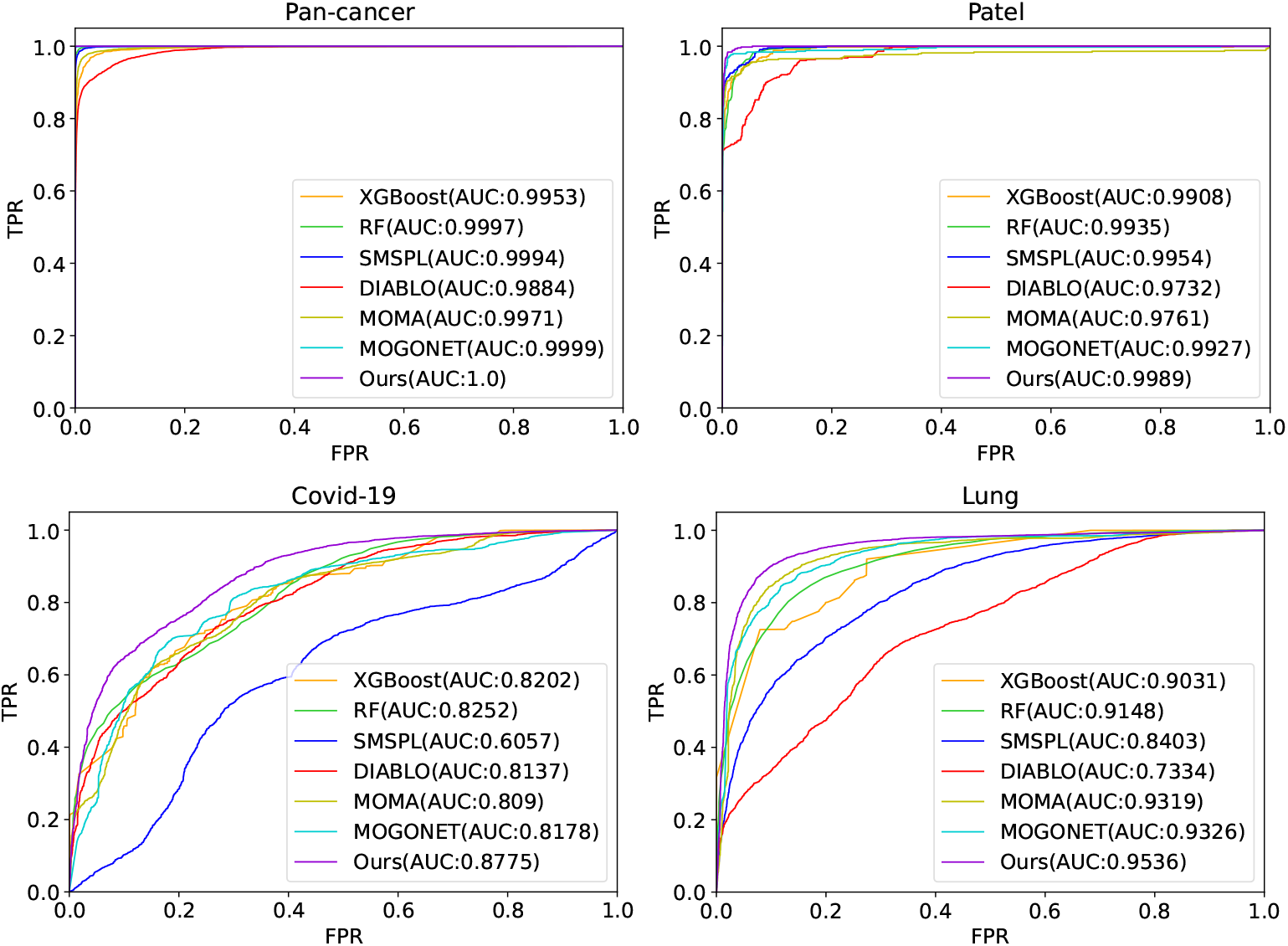
ROC curves for ISMI-VAE and baseline methods on four datasets, where the AUC of each method is shown in the legend.

The results in Fig. 3 are consistent with those in Table 1. On the one hand, the predictions of the methods are better on the Pan-cancer and Patel datasets than the other two datasets. On the other hand, the AUC of ISMI-VAE is greater than that of all other methods. This metric once again validates the excellent classification performance of ISMI-VAE.

### 4.4 ISMI-VAE has good interpretability

Under this experiment, we select the same number of important features in each modality based on the model’s importance score for the features and then concatenate them into a feature vector. We use a multilayer perceptron as a classifier to make predictions of diseases based on these features. Finally, we evaluate the interpretability of the model based on the performance of the classifier. ISMI-VAE measures the importance of features by attention weights, and its selection of features can be divided into two steps. First, the attention weights for the importance of gene expression data and the importance of SNV data are ranked from the largest to the smallest, and then, based on the ranking results, the important features of SNV data and gene expression data are selected in equal numbers for interpretability tests.

We selected different numbers of features for the experiment, The results of the comparison experiments are shown in Fig. 4. It can be seen from the figures that for the two simple tasks predicted within batches, ISMI-VAE selected features outperformed other methods for classification with 20 features. The performance gap between ISMI-VAE and the baseline method does not become smaller until the number of features becomes larger and the classification performance is saturated, which means that ISMI-VAE has good interpretability in the simple case. For the more difficult cross-batch prediction task, ISMI-VAE selects slightly worse features than the comparison method at 20 feature counts, and outperforms the comparison method in the more feature cases. This may be due to the fact that for the more difficult classification tasks, the deep learning models place more emphasis on the classification ability of the combined features.

**Fig. 4.**
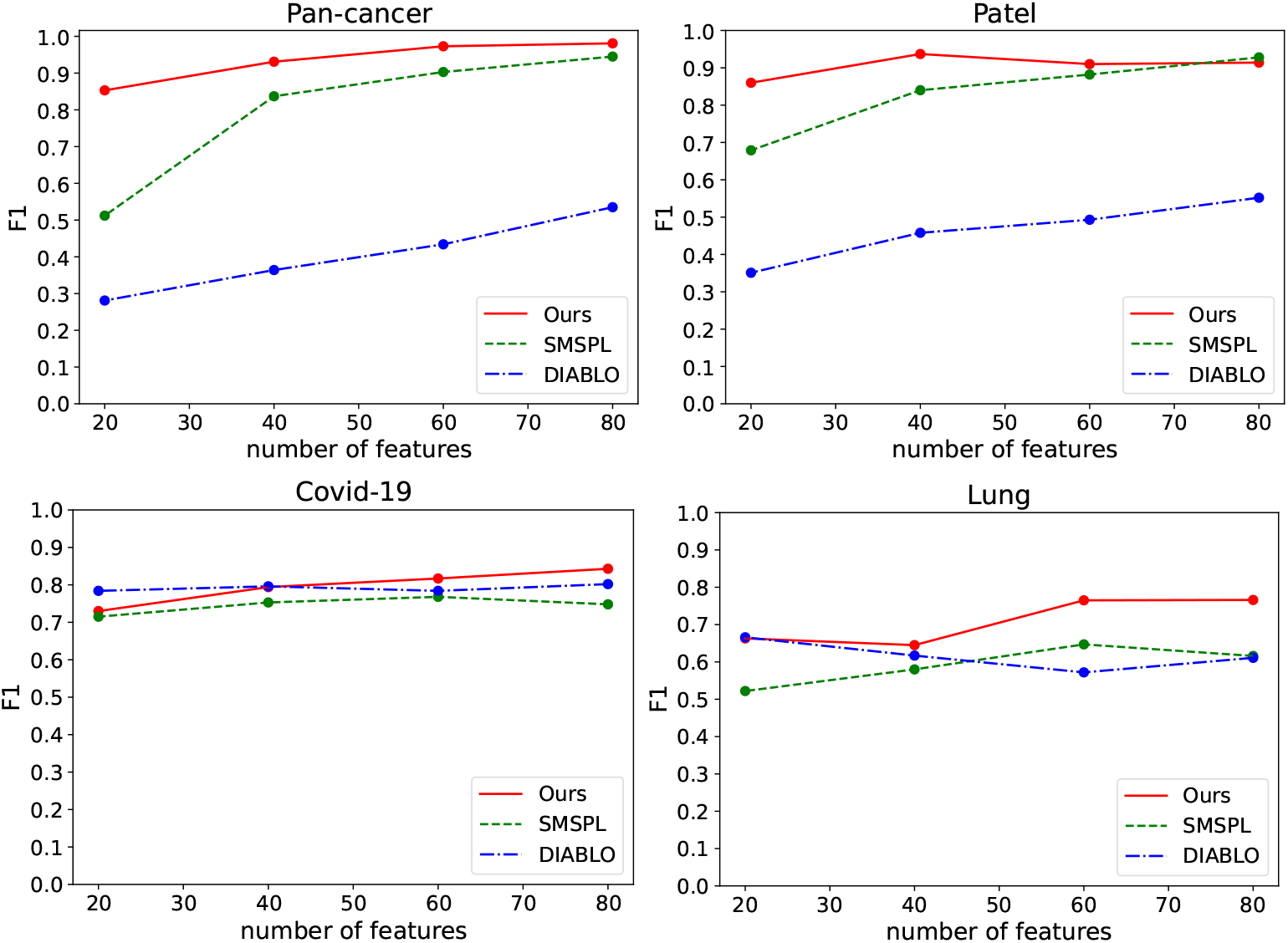
The classification results are based on 20, 40, 60, and 80 significant features, respectively, where different colors represent the significant features selected by different methods.

### 4.5 ISMI-VAE can find important SNVs

From the previous experimental analysis, it can be seen that the prediction of SNV data is significantly better than that of gene expression data in predicting the severity of Covid-19, which indicates that SNVs can cause the human body to have different resistance to Covid-19, while this aspect is relatively difficult to be reflected in gene expression. With the interpretability module of ISMI-VAE, we were able to further analyze the SNVs in the data that could determine the severity of Covid-19.

Fig. 5 shows the distribution of the attention weights of SNVs learned by the ISMI-VAE model. It can be seen that although the overall SNV attention weights are relatively high, the high weights are mainly concentrated in a few SNVs. Among these SNVs, there were only 100 SNVs with weights higher than 0.7, and we further analyzed these 100 SNVs. After excluding the missing values, we used the proportion of mutant reads as the status of this SNV and analyzed the difference between the status of SNVs on all severe Covid-19 cells and on all mild Covid-19 cells using rank sum test. The six SNVs with the highest level of significance of difference among these 100 SNVs and the distribution of their status on the two types of cells are shown in Figure 4.10.

**Fig. 5.**
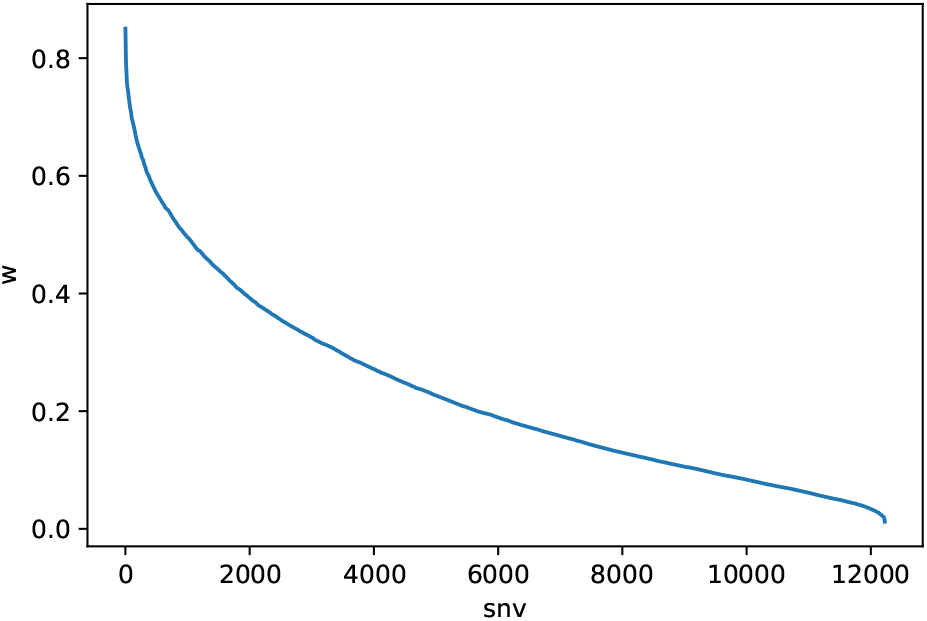
Attention weights of different SNVs in Covid-19 data determined by ISMI-VAE

As seen in Fig. 6, there is a clear difference in the SNV status in mild Covid-19 and severe Covid-19 cells, for example, at the locus chr22 29767537, the SNV status of severe Covid-19 cells tends more towards 1, while in mild Covid-19 cells, the SNV status at this locus tends more towards 0. In addition to this, the genes in which these loci are located have also been documented to be associated with Covid-19, such as TXNIP, the gene in which chr1 145993449 is located[24], S100A12, the gene in which chr1 153373787 is located[25], and UQCR10, the gene in which chr22 29767537 is located[26]. From the results of this experiment, we can see that ISMI-VAE has good interpretability, and relying on this interpretability, we are able to discover SNV loci or genes that affect the disease.

**Fig. 6.**
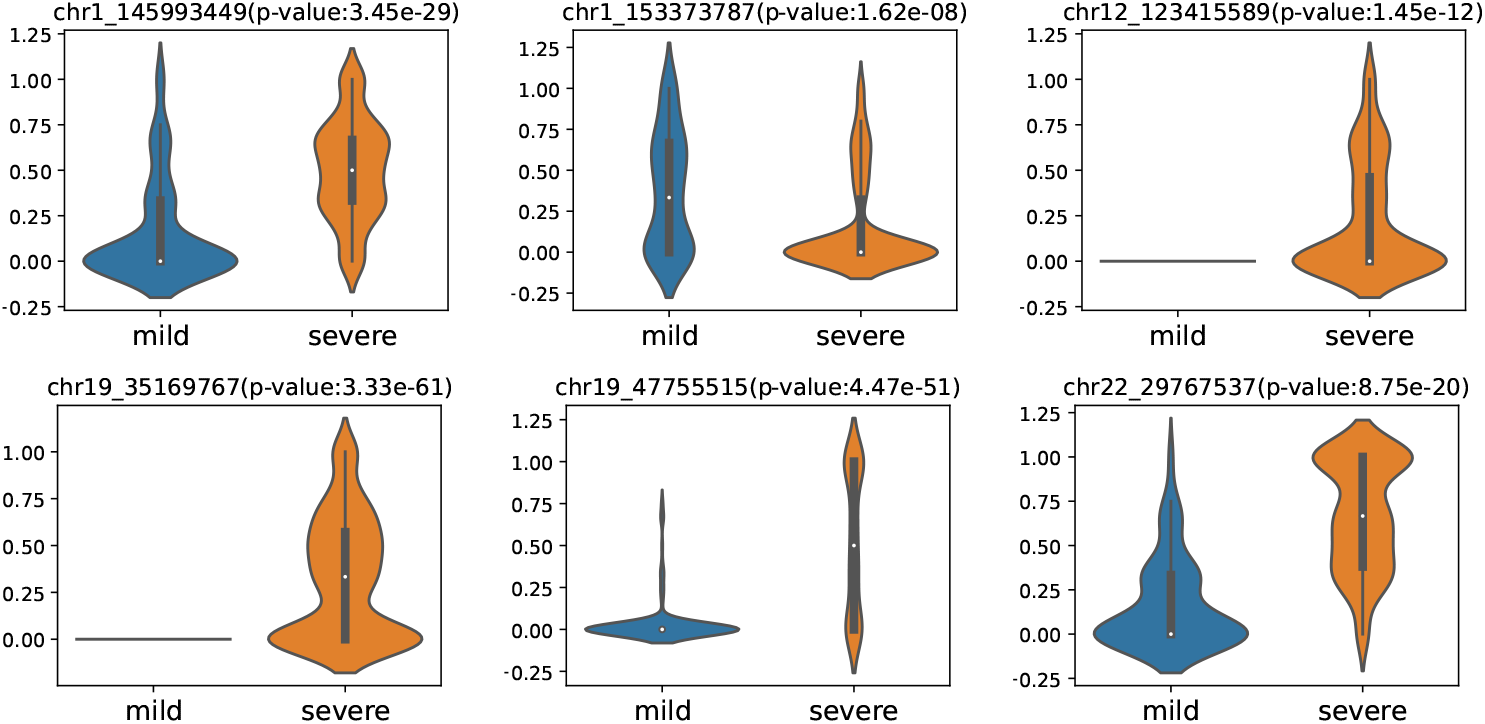
Significant SNVs in Covid-19 data analyzed by ISMI-VAE. The top 100 significant SNVs are first selected by weight ranking, and then the top 6 most significant SNVs are found by rank sum test. The figure caption shows the locus of this SNV as well as the p-value.

### 4.6 Parameter sensitivity analysis

The main parameter of ISMI-VAE is the regularization coefficient *λ*. Therefore, we analyze the sensitivity of this parameter. We measured the classification performance of the model when the parameter varied between 10^*−*4^ and 10. This classification performance is evaluated from two aspects. On the one hand, we test the overall classification performance of the model. On the other hand, we test the interpretability performance of the model, that is, we select the top 60 important features based on attention weight parameters, and then train them on a multilayer perceptron to evaluate their performance. The overall classification performance and interpretability performance are shown in Fig. 7 and Fig. 8.

**Fig. 7.**
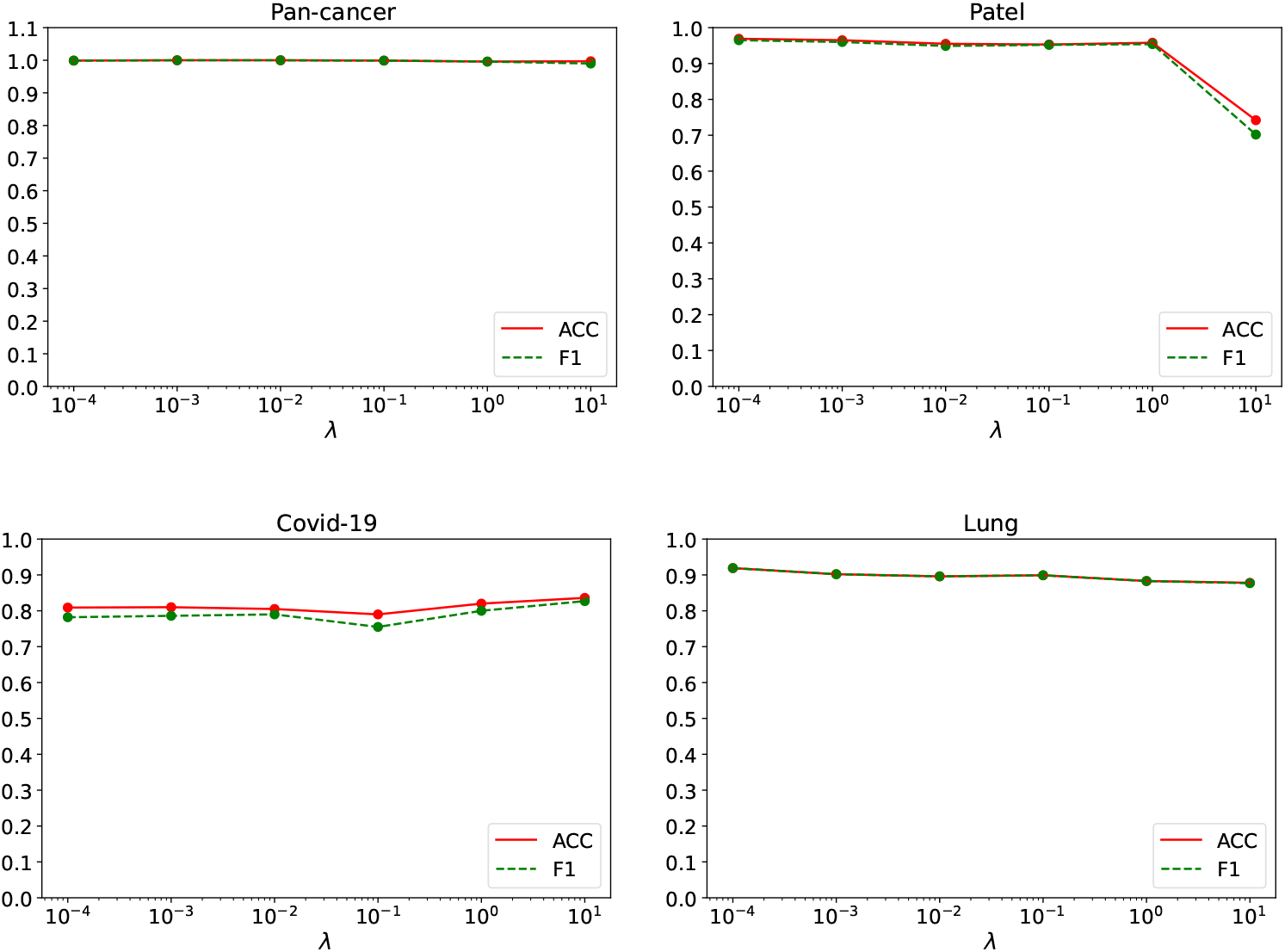
Classification performance of ISMI-VAE for different values of parameters *λ*.

**Fig. 8.**
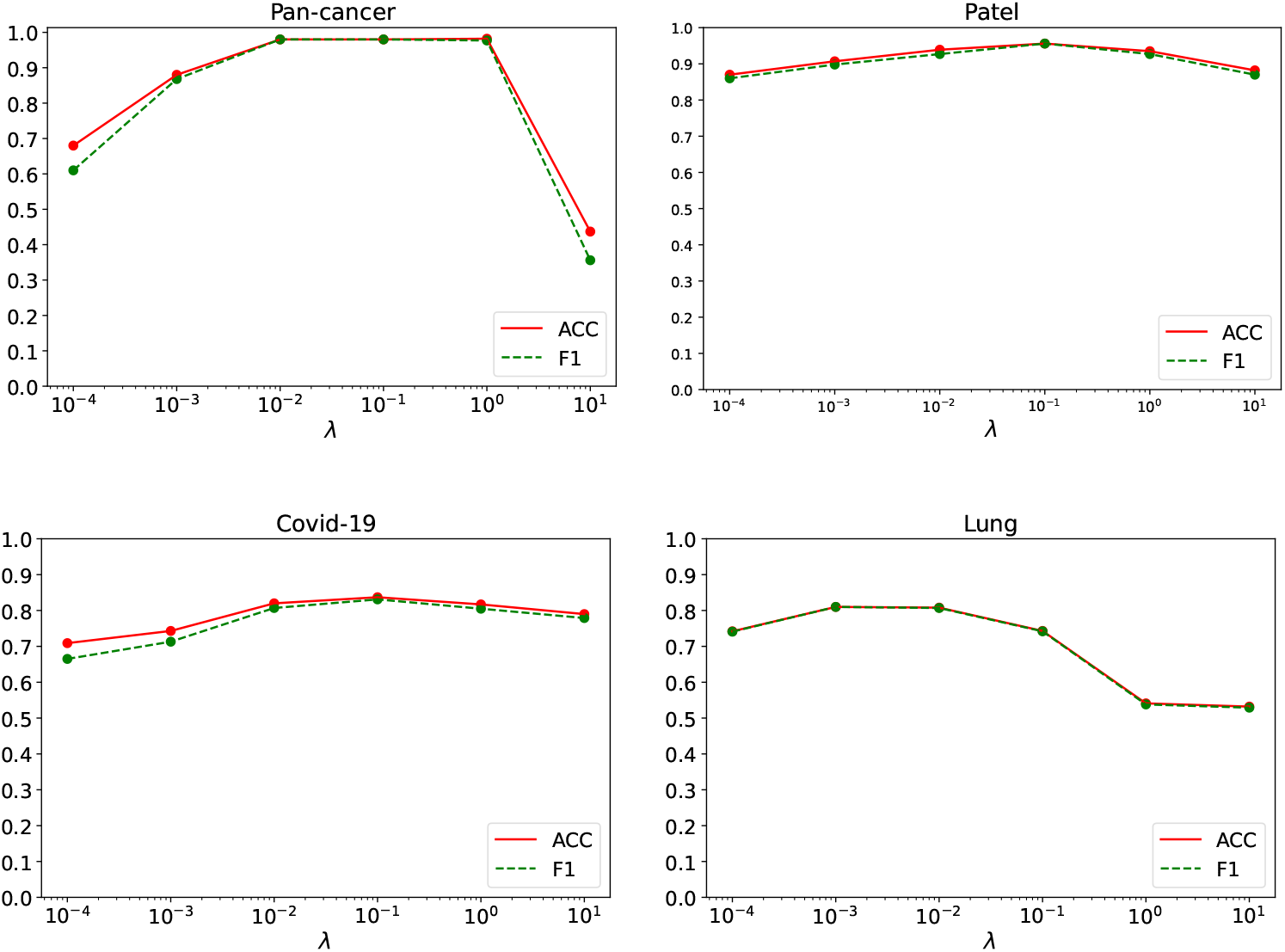
Interpretability of ISMI-VAE for different values of the parameter *λ*. The method for testing interpretability is the same as 4.4

As can be seen from the results, the overall performance is not greatly affected by *λ*, although there is a tendency for the overall performance to decrease slightly as *λ* increases, except for the Patel dataset where there is a relatively large decrease at *λ*=10. On the contrary, the performance on interpretability has a large fluctuation with the change of *λ*. Overall, the interpretable performance of the model decreases when *λ* is small. This may be due to the fact that the attentional regularization loss in the loss function loses its role when *λ* is too small. When *λ* is too large, the explainable performance of the model also shows a large decrease. This may be because the attentional regularization loss is too large, causing the model to focus only on the most important features and ignore other features that also possess more important information. Overall, *λ* values in the range of 1 to 10 are appropriate.

## 5 Conclusion

Compared to single-modal data, multimodal data can provide a more comprehensive biological insight due to the complementary information reflected between different data. With the development of biology, it has become easier to detect different modalities in single cells, and these data are commonly used, including gene expression, DNA methylation, etc. However, we found that SNV data, which are highly relevant to diseases, are rarely used in multimodal data integration. In this paper, we propose ISMI-VAE for integrating SNV data and gene expression data to classify disease cells.

ISMI-VAE draws on the idea of variational inference in VAE. It first constructs a latent variable model to describe the distribution of the two modal data and the interaction between them. Then, based on this latent variable model, ISMI-VAE constructs a neural network model to integrate the multimodal data and classify the disease cells. To further analyze the important genes associated with the disease and increase the interpretability of the model, we designed the latent variable model by introducing two latent variables to describe disease-causing and non-disease-causing genes. After that, we design an attention module on the neural network model to reflect the importance of different gene features by an attention vector trained simultaneously with the model, as well as for generating these two latent variables.

There is still room for improvement in ISMI-VAE. First, while the experiments in this paper demonstrate that SNV data can improve performance on some datasets, we find that it does not perform well on all datasets. What data are suitable for using SNV still needs further research. Secondly, ISMI-VAE was mainly designed for SNV data and gene expression data, and did not consider the integration of more modalities. However, this is not a big problem. This problem can be solved by modifying the structure of the latent variable model and adding appropriate inputs to the neural network model accordingly. This will be one of our future research works. Finally, in terms of interpretability, ISMI-VAE only considers the important features of individual modalities and does not address the connection between different modal data. In fact, it is very important to understand the interactions between different modal data. Therefore, how to use the interpretability module to analyze the interrelationship between different modal data is also something we need to further study in the future.

## 6 Methods

### 6.1 Generation of SNV data

We used gene expression data and SNV data for the experiments. Gene expression data were obtained by downloading directly from the data sources or by following the methods used by the authors who provided the data. Since these data sources do not consider or provide SNV data, we generated SNV data according to the standard process provided by Heaton[27]. The process can be divided into three steps. The first step is called remapping. We recreate the fastq file from the bam file and then map the reads to the reference sequence using minimap2[28, 29]. The second step uses freebayes[30] to call candidate variants. the third step uses vartrix to count cell allele counting. When these three steps are completed, two matrices are generated, an alternate matrix counting the number of mutated reads per locus and a reference matrix counting the number of unmutated reads per locus.

### 6.2 Latent-variable model

When designing the latent variable model, we took two main aspects into consideration. On the one hand, the features of the data can be decomposed into disease-related and unrelated components. For this reason, we introduced two latent variables to represent them separately. On the other hand, gene expression is also affected by SNVs, so the relationship between them needs to be considered when designing the latent variable model. In summary, the latent variable model we constructed is shown in Fig. 9. In the latent variable model, the variables *x*_*snv*_, *x*_*exp*_ are the SNV data and gene expression data, respectively, *y* is the sample label, *z*_1_ is the latent variable that is related to label *y* and *z*_2_ is the latent variable that is unrelated to label *y*.

**Fig. 9.**
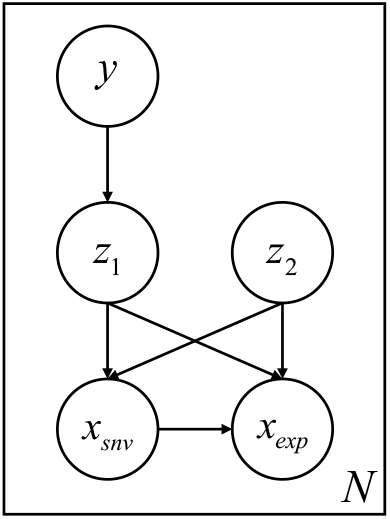
Latent-variable model

The generation process of the latent variable model can be described by the joint probability equation as:

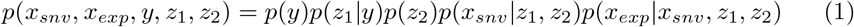

And in this process, each probability distribution is defined as:

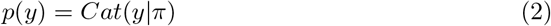

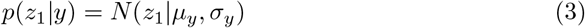

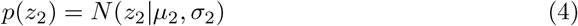

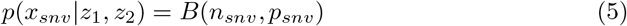

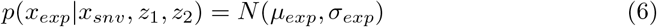

Where Cat is the category distribution, N is the Gaussian distribution, and B is the binomial distribution. For simplicity, let *x* = (*x*_*snv*_, *x*_*exp*_), *z* = (*z*_1_, *z*_2_). The objective of ISMI-VAE is maximizing the log-likelihood function log *p*(*x, y*).

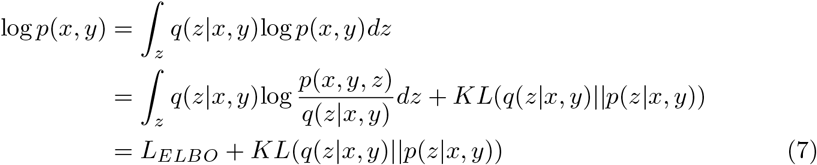

where *L*_*ELBO*_ is called the variational lower bound. Because *KL*(*q*(*z*|*x, y*)||*p*(*z*|*x, y*)) is always greater than or equal to 0, log *p*(*x, y*) ≥ *L*_*ELBO*_ and the log-likelihood function log *p*(*x, y*) can be maximized by maximizing *L*_*ELBO*_, which can be further represented as:

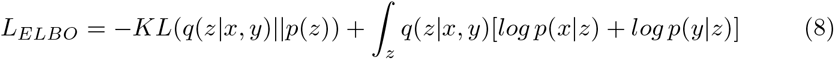

Since *x*_*exp*_ is influenced by *x*_*snv*_ and *z*_1_ and *z*_2_ are independent of each other, *L*_*ELBO*_ can be written as:

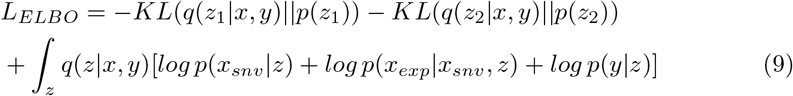

where *q*(*z* | *x, y*) and *p*(*x*_*exp*_ | *x*_*snv*_, *z*) are the Gaussian distribution, *p*(*x*_*snv*_ | *z*) is the binomial distribution and *p*(*z*) is standard Gaussian distribution. Their parameters are fitted by the neural network.

### 6.3 Neural Network Model

According to the latent variable model and loss function, the neural network model of ISMI-VAE is shown in Fig. 1. ISMI-VAE can be simply divided into feature selection module, modal fusion module, and prediction and reconstruction module. These modules are described below.

#### 6.3.1 Feature Selection

First, ISMI-VAE determines the importance of different features in the input data *x*_*snv*_ and *x*_*exp*_ through a feature selection module and divides the input data into two parts according to the importance of the features. This module is mainly used to achieve interpretability of the whole model, and ISMI-VAE achieves this by the attention mechanism as follows:

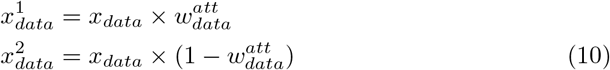

where 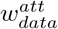 is the attention weight of the corresponding input data, which is a learnable one-dimensional vector with the same length as the number of features of the input data and taking values in the interval [0,1]. The larger the value on this attention vector, the more important the corresponding feature is for model classification. 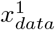 is the part of the input data *data* that is relevant to the label *y*, i.e., the important features. 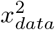 is the part of the input data data that is not relevant to the label *y*, i.e., the unimportant features. The value range of *data* is *exp, snv*_*a*_, *snv*_*r*_, *x*_*exp*_ is gene expression data, 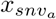 is the alternate matrix of SNV data, 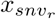 is the reference matrix of SNV data. 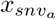 and 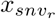 share the same parameters.

To ensure that every value in 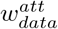 is in the interval [0,1], a trick is used. ISMI-VAE first generates a random attention vector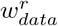. and then uses the Sigmoid function to restrict the range of values of 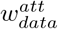:

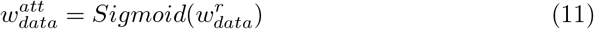

Considering that we expect to select features with 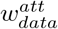, it is better to have 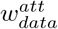 equal only to 0 or 1 to represent selecting a feature or not selecting a feature, respectively. Therefore, in this paper, the attention vector is further improved. Specifically, ISMI-VAE uses Gumbel Softmax[31] to sample the attention vector.

#### 6.3.2 Modal Fusion

The modal fusion module is used to fuse the multimodal information obtained from the previous step of ISMI-VAE and generate latent variables *z* based on the fused information. We designed two modal fusion modules with identical structures but different parameters to generate latent variables *z*_1_ and *z*_2_, respectively, Firstly, ISMI-VAE generates intermediate variables as follows:

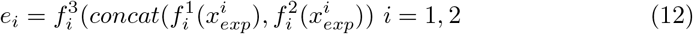

where *i* = 1 represents the fusion of important features of two modalities, *i* = 2 represents the fusion of unimportant features of two modalities. 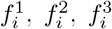 are the corresponding multilayer perceptron. *Concat* represents concatenation of two vectors. Then, the ISMI-VAE uses the multilayer perceptron to predict the parameters of the posterior probability *q*(*z*_*i*_|*x, y*) according to the intermediate variable *e*_*i*_ as follow:

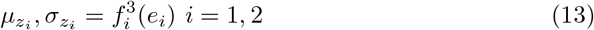

If the latent variable *z*_*i*_ is sampled directly according to posterior probability *q*(*z*_*i*_ | *x, y*), the whole model will not be derivable. Therefore, ISMI-VAE uses the same solution as VAE, that is, ISMI-VAE samples the latent variable z by reparameterization:

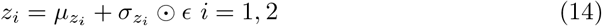

#### 6.3.3 Rediction and Reconstruction

In the prediction and reconstruction module, ISMI-VAE predicts label *y* based on latent variable *z*_1_, and reconstruct sample *x* based on latent variables *z*_1_ and *z*_2_. ISMI-VAE uses multilayer perceptron *f* ^5^ to predict label *y*, multilayer perceptron *f*^*snv*^ to reconstruct SNV data, and multilayer perceptron *f*^*exp*^ to reconstruct gene expression data:

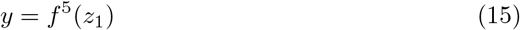

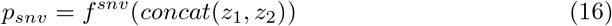

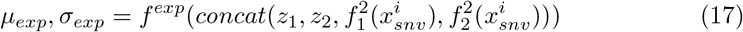

In summary, the loss function of the model is as follows:

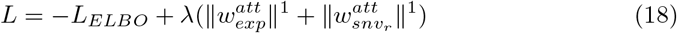

where *L*_*ELBO*_ is the variational lower bound loss, ∥·∥^1^ represents l1 regularization, and *λ* is the weight coefficient for weighing feature selection and variational lower bound optimization, which is used to control the number of important feature selections. The larger *λ* is, the more ISMI-VAE tends to assign larger weights to the few features. Only 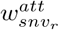 is regularized here because 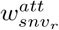 and 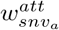 share parameters. By optimizing the variational lower bound loss, the model can learn the latent variables of the data, and through regularization constraints, the model can force itself to select only the necessary features to generate the latent variable *z*, and predict the label *y*, By optimizing the variational lower bound loss, the model can learn the correct latent variable distribution, and through regularization constraints, the model can force itself to select only the necessary features to generate the latent variable *z*_1_ and *y*.

## Acknowledgments

We appreciate the help and advice of our partners and reviewers.

## Declarations

### Availability of data and materials

The datasets analyzed in this study are available from the Gene Expression Omnibus (GEO) repository under the following accession numbers:GSE57872[20],GSE157220[21],GSE149689[12], GSE123902[22].

### Ethics approval

### Ethics approval and consent to participate

Not applicable.

